# An Updated Gene Regulatory Network reconstruction of multidrug-resistant *Pseudomonas aeruginosa* CCBH4851

**DOI:** 10.1101/2022.05.12.491685

**Authors:** Márcia da Silva Chagas, Fernando Medeiros Filho, Marcelo Trindade dos Santos, Marcio Argollo de Menezes, Ana Paula D’Alincourt Carvalho-Assef, Fabricio Alves Barbosa da Silva

## Abstract

**BACKGROUND:** Healthcare-associated infections due to multidrug-resistant (MDR) bacteria such as *Pseudomonas aeruginosa* are significant public health issues worldwide. A system biology approach can help understand bacterial behavior and provide novel ways to identify potential therapeutic targets and the development of new drugs. Gene regulatory networks (GRN) are an example of interaction representation *in silico* between regulatory genes and their targets.

**OBJECTIVES:** In this work, we update the reconstruction of the MDR *P. aeruginosa* CCBH4851 GRN, and analyze and discuss its structural properties.

**METHODS:** We based this study on the gene orthology inference methodology using the reciprocal best hit method. The *P. aeruginosa* CCBH4851 genome and GRN, published in 2019, and the *P. aeruginosa* PAO1 GRN, published in 2020, were used for this update reconstruction process.

**FINDINGS:** Our result is a GRN with a larger number of regulatory genes, target genes, and interactions compared to the previous networks, and its structural properties are consistent with the complexity of biological networks and the biological features of *P. aeruginosa*.

**MAIN CONCLUSIONS:** Here, we present the largest and most complete version of *P. aeruginosa* GRN published to this date, to the best of our knowledge.

## INTRODUCTION

*Pseudomonas aeruginosa* is a ubiquitous and opportunistic pathogen of which infections can affect the lower respiratory tract, skin, urinary tract, eyes, soft tissues, surgical wound, gastrointestinal system, among others, leading to bacteremia, endocarditis, and other complications, particularly in health care settings and in immunocompromised patients.^(1-3)^ Moreover, this Gram-negative bacteria is one of the most difficult to treat,^(4)^ due to its intrinsic resistance, acquisition of resistance through chromosomal gene mutations, and horizontally acquired resistance mechanisms to multiple drugs.^(3)^ Multidrug resistance (MDR) imposes the central difficulty in the selection of appropriate antibiotic treatment and reduces treatment options, especially in nosocomial settings such as healthcare-associated infections (HAI).^(5,6)^ HAI is a severe public health issue related to high rates of morbidity and mortality in hospitalized patients and excessive healthcare costs.^(7)^ Worldwide, *P. aeruginosa* is one of the most prevalent agents of HAI.^(8)^

In Brazil, the Brazilian Health Surveillance Agency ^(9)^ ranked *P. aeruginosa* as the third most common causative agent of HAI in hospitalized patients in adult intensive care units (ICU) and the second in pediatric ICU, being nearly 40% of the reported strains resistant to carbapenems ^(9)^. This class of beta-lactam antibiotic has been widely administered worldwide for treating *P. aeruginosa* infections and other MDR Gram-negative bacterial infections.^(10)^ Indeed, a significantly higher mortality rate was observed among patients infected with MDR *P. aeruginosa* clones (44.6%) compared to those infected with non-MDR (24.8%).^(6)^

The most epidemiologically important mechanism of carbapenem resistance is the production of carbapenemases. Among MDR P. aeruginosa clinical isolates in Brazil, the most prevalent carbapenemase is the São Paulo metallo-β-lactamase (SPM-1).^(11)^ This enzyme is encoded by the gene *blaSPM-1*, located on the *P. aeruginosa* chromosome,^(12)^ and it confers resistance to almost all classes of beta-lactams. The first register of an MDR *P. aeruginosa* strain carrying the *blaSPM-1* gene found in Brazil is from 2003.^(13)^ Widely disseminated in distinct Brazilian geographic regions, SPM-1-producing *P. aeruginosa* is associated with the clone SP/ST277 and has been isolated from hospital sewage systems, rivers, and microbiota of migratory birds.^(11,12)^ The strain *P. aeruginosa* CCBH4851, in which this article is based on, belongs to clone SP/ST277, and was involved in an endemic outbreak in Brazil in 2008.^(14)^ This strain is resistant to most antimicrobials of clinical importance, such as aztreonam, amikacin, gentamicin, ceftazidime, cefepime, ciprofloxacin, imipenem, meropenem, and piperacillin-tazobactam, being susceptible only to polymyxin B, and has several mechanisms of mobile genetic elements.^(2, 14)^

To better understand *P. aeruginosa*’s behavior, a more comprehensive knowledge of gene expression patterns predicted by the analysis of its gene regulatory network (GRN) is of great value. A GRN consists of a set of transcription factors (TF) that interact selectively and nonlinearly with each other and with other molecules in the cell to regulate mRNA and protein expression levels.^(15)^

Mathematical modeling and computational simulations are approaches for analyzing the GRN and other complex cellular systems influenced by numerous factors. These models allow the construction of biological networks, predict its behavior under unusual conditions, identify how a disease might develop, and intervene in such development to prohibit cells from reaching undesirable states.^(16)^ In addition, due to their lower cost and high accuracy, such approaches contribute to developing new drugs. ^(17)^

The *P. aeruginosa* PAO1 strain had its genome sequence published in 2000, providing information regarding genome size, genetic complexity, and ecological versatility.^(18)^ It has been extensively studied since then. Published in 2011 by Galán-Vasquez *et al*.,^(19)^ the first *P. aeruginosa* GRN was based on the PAO1 strain (PAO1-2011). In 2019, Medeiros *et al*.^(2)^ described a GRN reconstruction of the CCBH4851 strain (CCBH-2019). Finally, in 2020, Galán-Vasquez *et al*.^(20)^ published the updated GRN of *P. aeruginosa* with the PAO1 strain (PAO1-2020), which was much larger than the previous ones, containing new interactions. All works analyzed the GRNs main structural properties and regulatory interactions.

This manuscript describes (CCBH-2022), an updated GRN of the MDR *P. aeruginosa* based on the CCBH4851 strain, using as references both CCBH-2019 and PAO1-2020. We characterize regulators, target genes (TGs), transcription factors (TFs), auto-activation interactions and influential genes of the network.

We analyze the main structural properties of the network, such as degree distribution, clustering coefficient and relative abundance of network motifs. We compare results of our analyses with those from previous GRNs.

## MATERIALS AND METHODS

In this work we study the *P. aeruginosa* CCBH4851 strain, which is deposited at the Culture Collection of Hospital-Acquired Bacteria (CCBH) located at the Laboratório de Pesquisa em Infecção Hospitalar, Instituto Oswaldo Cruz/Fundação Oswaldo Cruz (Fiocruz) (WDCM947; 39 CGEN022/2010). The genome sequence is available in the GenBank database (Accession CP021380.2).^(14)^

CCBH-2019 and PAO1-2020 were the bases for the reconstruction of this GRN. CCBH-2022 results from the orthology analysis between the *P. aeruginosa* PAO1 and CCBH4851 gene sequences. The CCBH-2022 model also inherits the orthologs between CCBH4851 and *P. aeruginosa* PA7 ^(21)^ and *P. aeruginosa* PA14 ^(22)^ strains, which were already present in CCBH-2019.

The evolutionary histories of genes and species reconstruction are based critically on the accurate identification of orthologs.^(23)^ Orthology refers to a specific relationship between homologous characters that arose by speciation at their most recent point of origin,^(24,25)^ a common ancestor.

One of the most common approaches to determine orthology in comparative genomics is the Reciprocal Best Hits (RBH), which BLAST relies on.^(26)^ An RBH occurs when two genes, each in a different genome, find themselves as the best scoring match in the opposite genome.^(27, 28)^

Regulatory interactions between TFs and TGs in the PAO1 GRN were propagated to the CCBH-2022 GRN if the TF and the TG formed a RBH. An algorithm was designed and implemented by Medeiros *et al*.^(2)^ using the Python programming language to automate and generate a list of RBHs in a tabular format (available as Supplementary Data). All the protein sequences from *P. aeruginosa* CCBH4851 (P1) and *P. aeruginosa* PAO1 (P2) were considered. BLAST+ ^(29)^ were used to query the proteins from P1 against those from P2 (forward results) and P2 against P1 (reverse results). Each P1’s query sequence was considered in turns, and its best match from P2 was identified from forwarding results (x). Likewise, each P2’s query sequence was considered from the reverse results, with its best match in P1 (x’). If x = x’, then they are RBH. Local BLASTP searches of each protein set against the other were executed, with the following cut-off parameters: *identity* ≥ 90%, *coverage* ≥ 90%, and *E-value* ≤ 1 e-5, showing the results in tabular format.

If the search returned no hits, the gene was considered to have no ortholog within the opposite genome. Manual BLASTP was used to prevent false negatives, aligning these gene sequences with the opposite genome, considering the above parameters. If they still returned no hits but were present in either PAO1-2020 or CCBH-2019, the results were evaluated with a literature search to determine if they were accurate and whether they should or not be part of the CCBH-2022.The final GRN table is available as Supplementary data and is organized into six columns: Regulatory gene, Ortholog of the regulatory gene, Target gene, Ortholog of the target gene, Mode of regulation, and Reference. The first column lists the regulatory genes of *P. aeruginosa* CCBH4851 while the second column contains orthologs of regulatory genes in the reference strain (PAO1 and PA7 or PA14 from the exclusive interactions in CCBH-2019; the same applies to TG’s orthologs). The third column refers to the target gene in *P. aeruginosa* CCBH4851 while the fourth column lists orthologs of TGs in the reference strain. The fifth column describes the mode of regulation and the sixth column indicates the source of the corresponding data.

The set of interactions between transcription factor proteins and the genes that they regulate in an organism define a directed graph. For the computational analysis, the structure of GRN can be represented as a directed graph, formed by a set of vertices (or nodes) connected by a set of directed edges (or links). Basic network measurements are related to vertex connectivity, occurrence of cycles, and the distances between pairs of nodes, among other possibilities.^(30)^

The degree of vertices is the most elementary characterization of a node, being the k(i) defined as its number of edges. In directed networks, there are incoming (k-in degree) edges and outgoing (k-out degree) edges.^(31)^ The degree distribution can follow a functional form P(k) = Ak ^-γ^, called power-law distribution, where P(k) is the likelihood that a randomly chosen node from the network has k direct interactions, A is a constant that ensures that the P(k) values add up to one, and γ is the degree exponente.^(32-36)^ According to Albert (2005),^(37)^ this function indicates high diversity of node degrees, with the P(k) value decaying as a power law that is free of a characteristic scale, resulting in the absence of a typical node in the network that could be used to characterize the rest of the nodes. Most real networks with structural information available exhibit this scale-free behavior, deviating from a Poisson distribution expected in a classical random network.^(38,39)^

Studies have shown a scale-free structure in cellular metabolic networks,^(32,40)^ protein interaction networks, including in câncer,^(41,42)^ transcription regulatory networks, and GRN.^(20,43-45)^

Following the literature,^(36,37,46-48)^ there is some qualitative and quantitative characteristics to ensure that a network is scale-free: the power-law distribution appears as a straight line on a logarithmic plot; the γ value usually is in the range 2<γ<3; and the presence of high-degree nodes, called hubs, dominating the network, with most nodes clustered around them.

The hubs show the absence of a uniform connectivity distribution in the network, presenting the 70-30 rule (also referred to as the Pareto principle), with small-degree nodes being the most abundant. However, the frequency of high-degree nodes decreases slowly.^(37)^ Hubs are fundamental for determining therapeutic targets against an infectious agente.^(2)^

Scale-free networks are heterogeneous,^(49)^ so random node disruptions (the 70%) do not lead to a significant loss of connectivity. However, the loss of the hubs (the 30%) causes the breakdown of the network into isolated clusters.^(50)^ Some studies validate these general conclusions for cellular networks.^(51-53)^

In the GRN, determining the vertices with the highest k-out degrees is a method for identifying a hub,^(2)^ The degree threshold is the exact number of interactions that characterize a hub, and this criteria differs from a study to another.^(54)^ The degree threshold adopted in this work was the average number of connections of all nodes having at least two edges, resulting in a cut-off value of 16 connections. Motifs are patterns of connectivity, a small set of recurring regulation patterns from which the networks are built ^(55,56)^ that are associated with specific functions.^(57)^ A triangle, i.e., three fully connected vertices, is the minimum of a motif.^(47)^ These genes are a regulator, X, which regulates Y, and gene Z, which is regulated by both X and Y.^(58)^ Triangles can be closed (three connections within the set) or open (two edges).^(38)^ This 3-genes motif is the feedforward loop (FFL), and the most common in GRN, appearing in gene systems in bacteria and other organisms,^(59,60)^ with the possibility of either activation and repression in each of the three regulatory interactions.^(61)^ The coherent type-1 FFL and the incoherent type-1 FFL occur more frequently in transcriptional networks.^(58)^

The clustering coefficient is the probability that two genes with a common neighbor in a graph are also interconnected.^(19)^ There are two popular definitions of this measure: the local and global clustering coefficient. The local clustering coefficient of vertex i, C_i_, is defined as C_i_ = 2e_i_ / k_i_ (k_i_ - 1), where e_i_ is the number of edges connecting node i with degree k_i_, and k_i_ (k_i_ - 1) / 2 is the maximum number of edges in the neighborhood of node i.^(36)^ In GRNs, the local clustering coefficient C(i) is interpreted as the interaction between genes forming the regulatory groups.^(2)^

The clustering coefficient of a network, C, is calculated by the average of C_i_ over all vertices.^(19,62)^

Not considering the directionality of the edges, the global clustering coefficient is the ratio between the number of closed triangles and the total number of triangles (open or closed) in the network.^(2)^

C(k) represents the mean clustering coefficient over the vertices with degree k.^(36)^ Some biological networks tend to present high clustering coefficient values.^(47)^ The network density measure is the number of edges of the network over the maximum possible number of edges, measuring the interconnectivity between vertices, and is strongly correlated to the potential to generate gene expression heterogeneity.^(63)^

The network diameter is the path length between the two most distant nodes.^(36)^ The average path length is the measure that indicates the distances between pairs of vertices (the average of the shortest path length over all pairs of nodes in the network).^(46)^

Several genes are connected with one another in the GRN. When the nodes interact through a direct or an indirect link (intermediate connections), they are considered as a part of a connected component. These associations are the concept of network connectivity, and for this analysis in the present work, network interactions were considered undirected.^(2)^

Analyzing the structural characteristics (connected components, hubs, and motifs) can help determine the best approach to disturb a network to promote a desired phenotype in the cell.^(64)^

For the CCBH-2022 structural analyses, the R programming language and RStudio were used.^(65)^ Scales, dplyr, tibble, readr and igraph packages were used for data manipulation and plotting of the structural analyses.^(2,20,66)^ The igraph library was used to compute most of the properties described above: the in and out degrees, centrality, clustering coefficients, feed-forward loop motifs, connectivity, cycles, paths, and hierarchical levels analyses.^(67)^

The illustrations of the GRN, the hub’s network and the connectivity analysis were made in Cytoscape.^(68)^ All figures are presented with higher resolution in the Supplementary Material.

The Supplementary Data, the codes for the structural analysis in R and for finding RBH in python, implemented by Medeiros *et al*.^(2)^, and the CCBH-2022 file in CSV format are available in our Github repository (https://github.com/FioSysBio/CCBH2022).

## RESULTS

The CCBH-2022 consists of 5507 regulatory interactions among 3291 gene products, of which 217 were identified as regulatory genes and 3071 as target genes. Of the 217 regulatory proteins, 87 are TFs, 19 are sigma factors (SF), and 13 are RNAs. Of these 13 RNAs, 11 are SF as well. The tables containing their relations are present in the Supplementary Material.

Given the 6577 predicted protein-coding genes of *P. aeruginosa* CCBH4851, the model organism in this study, the current network represents roughly 50% of the genome, against 16.52% from CCBH-2019.

Specific regulatory genes and their interactions were kept as described in the CCBH-2019, such as the ones resulting from the *P. aeruginosa* PA7 and *P. aeruginosa* PA14 orthology, and in dedicated biological databases and scientific literature, e.g., IHF (integration host factor). This bacterial DNA-bending protein, essential in gene expression regulation, is absent in the CCBH4851 genome. However Delic-Attree *et al*. (1995)^(69)^ demonstrated that *P. aeruginosa* contains the IHF protein composed of the products of the *himA* and *himD* genes. These genes act in combination as a TF for several TGs,^(2)^ and all were listed as regulatory genes in CCBH-2019. Consequently, equivalent notations to the previous CCBH4851 GRN were maintained.

CCBH-2022 has 5507 edges, and these interactions were classified into activation (“+”), repression (“-”), dual (“d”, when, depending on the conditions, the regulatory gene act as an activator or a repressor), and unknown (“?”). An illustration of CCBH-2022 is presented in Fig.1.

**Fig. 1:**
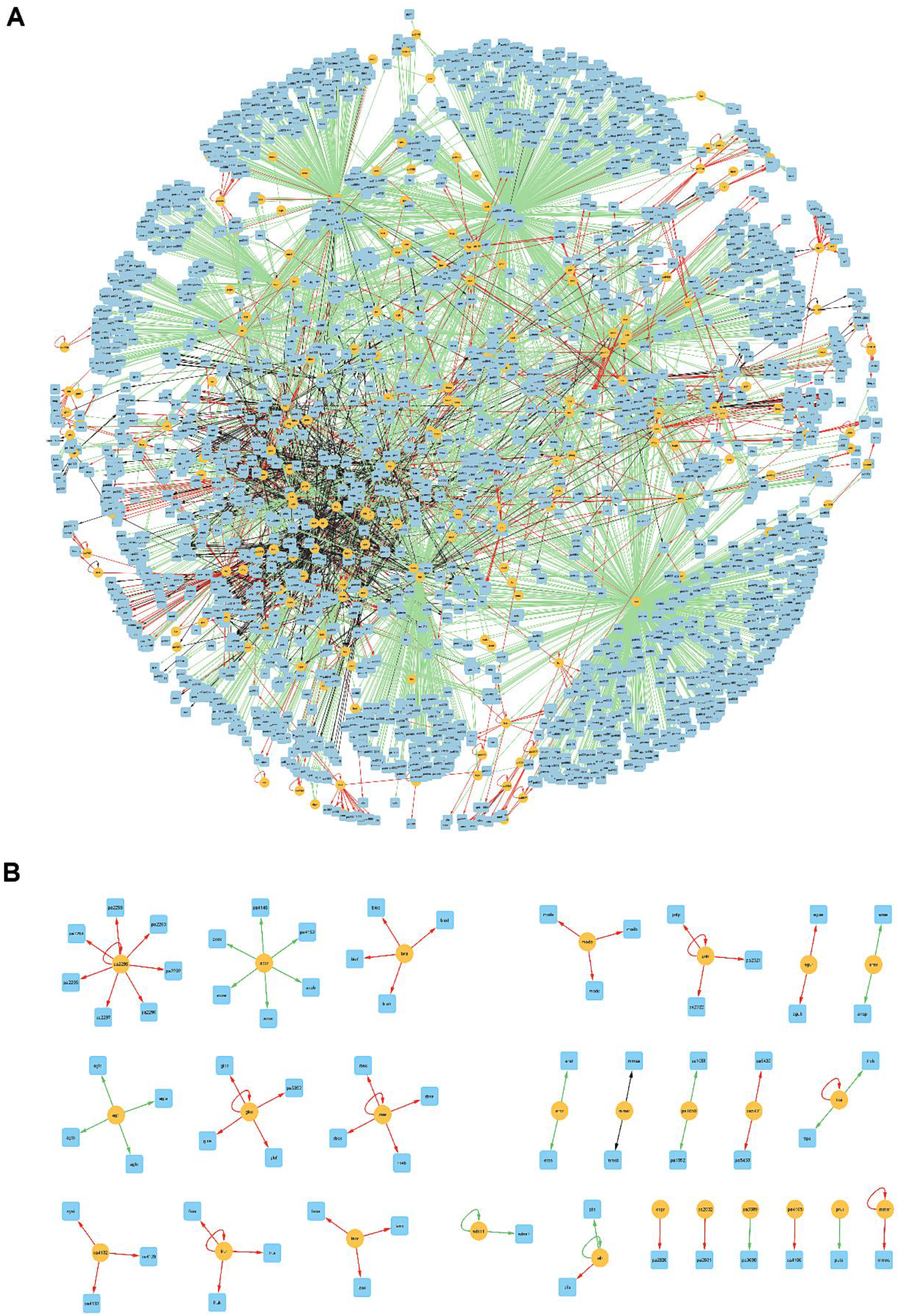
visualization of the CCBH-2022. Yellow circles indicate regulatory genes, light blue circles indicate target genes, black lines indicate an unknown mode of regulation, green lines indicate activation, and red lines indicate repression.Purple lines indicate a dual-mode of regulation. A: the GRN large highly connected network component; B: all regulatory and TGs with no connections with A.

Regarding the structural measurements of the updated network, the summarized statistical results are presented in Table 1. It contains the standard measures (the number of nodes and edges, number of autoregulatory motifs, network diameter, and average path length), the number of feed-forward motifs, and clustering coefficients. Also, Table I presents a comparison with data from the PAO1-2011, CCBH-2019, PAO1-2020, and CCBH-2022.

**TABLE 1.** Comparison of structural statistic measures between the PAO1-2011, CCBH-2019, PAO1-2020, CCBH-2022

|  | PAO1-2011 | CCBH-2019 | PAO1-2020 | CCBH-2022 |
| --- | --- | --- | --- | --- |
| Vertices | 690 | 1046 | 3006 | 3291 |
| Edges | 1020 | 1576 | 5040 | 5507 |
| Regulatory genes | 76 | 138 | 173 | 217 |
| Target genes | 593 | 908 | 2709 | 3071 |
| Positive regulation | 779 | 772 | 3851 | 3873 |
| Negative regulation | 218 | 454 | 390 | 661 |
| Dual regulation | 11 | 13 | 10 | 16 |
| Unknown regulation | 12 | 337 | 789 | 957 |
| Autoregulation (total) | 29 | 72 | 50 | 93 |
| Positive autoregulation | 16 | 21 | 24 | 30 |
| Negative autoregulation | 13 | 39 | 15 | 46 |
| Unknown autoregulation | - | 12 | 11 | 17 |
| Feed-forward loop motifs (total) $\alpha$ | 137 | 208 | 702 | 951 |
| Coherent type I feed-forward loop motifs $\alpha$ | 82 | 79 | 226 | 240 |
| Incoherent type II feed-forward loop motifs $\alpha$ | 3 | 4 | 8 | 10 |
| Density | 2.12e-03 | 1.44e-03 | 6.07e-04 | 5.68e-04 |
| Diameter | 9 | 12 | 12 | 12 |
| Average shortest path length | 04.08 | 4.80 | 04.01 | 4.65 |
| Global clustering coefficient | 2.28e-02 | 3.2e-02 | 3.03e-03 | 4.27e-03 |
| Local clustering coefficient | 2.5e-01 | 1.92e-01 | 1.63e-01 | 1.89e-01 |

$\alpha$ : number of feed-forward loop motifs determined using the igraph package.

The CCBH-2022 had a density of 5.68e-04, much higher than PAO1-2011 (2.12e-03) and CCBH-2019’s (1.44e-03) densities. It was slightly lower than the density of PAO1-2020 (6.07e-04) but showed the same order of magnitude. The diameter was 12, the same as the CCBH-2019 and PAO1-2020 and higher than PAO1-2011, which was 9.

The average shortest path distance was 4.65, higher than PAO1-2020 (4.01) but slightly lower than CCBH-2019 (4.80).

The degree distributions of the four networks can be seen in Fig. 2 (A, B, C, D), with A and B being the incoming and C and D the outcoming degree distribution. Fig.2B, D is on a double logarithmic axis and the straight line is consistent with a power-law distribution. For k-in, the estimated value for γ was 2.79, within the range 2<γ<3, consistent with a power law distribution. For the PAO1-2020 GRN the corresponding value was 2.67, being 2.89 for the CCBH-2019 GRN and 2.71 for the PAO1-2011 GRN.

**Fig. 2:**
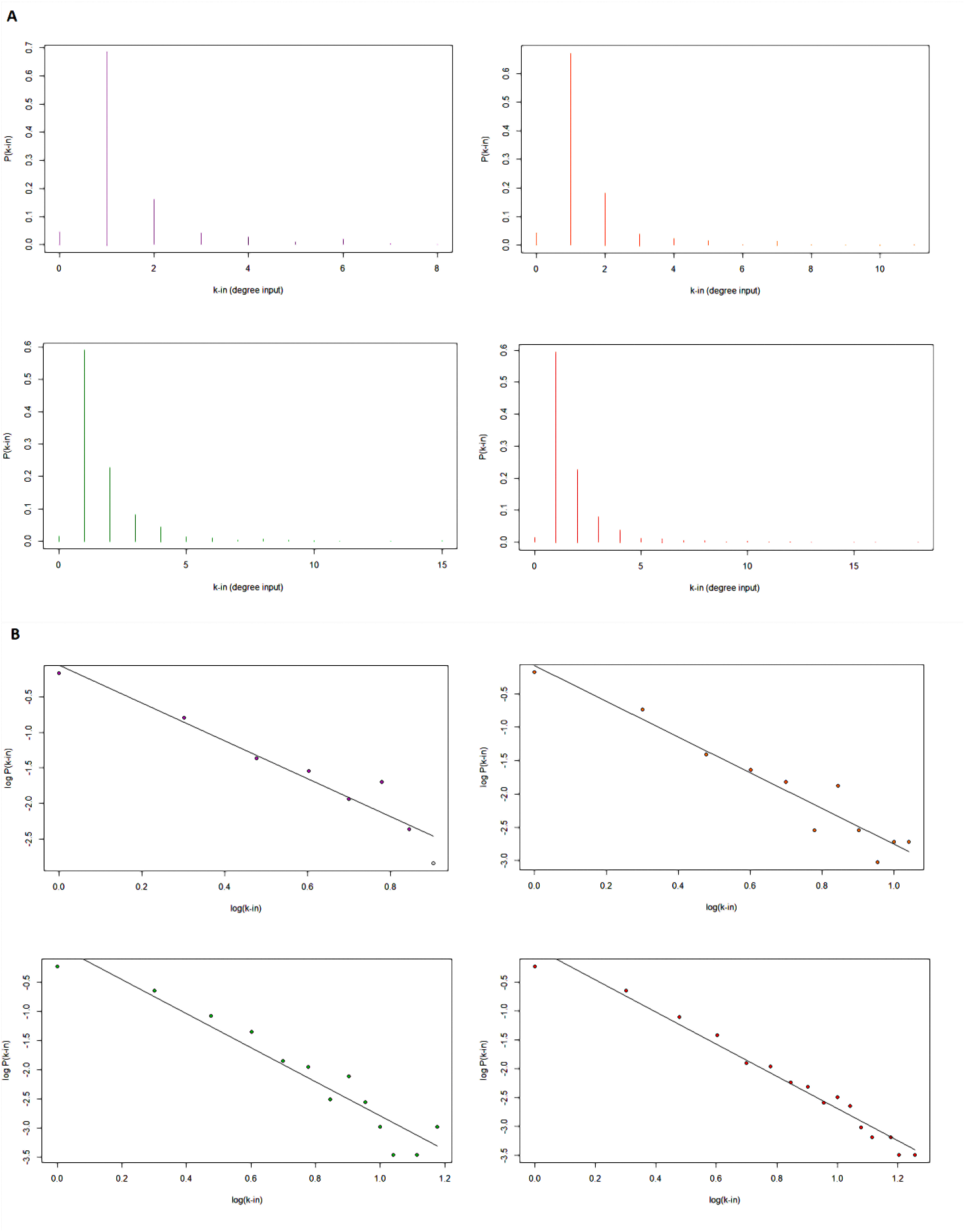
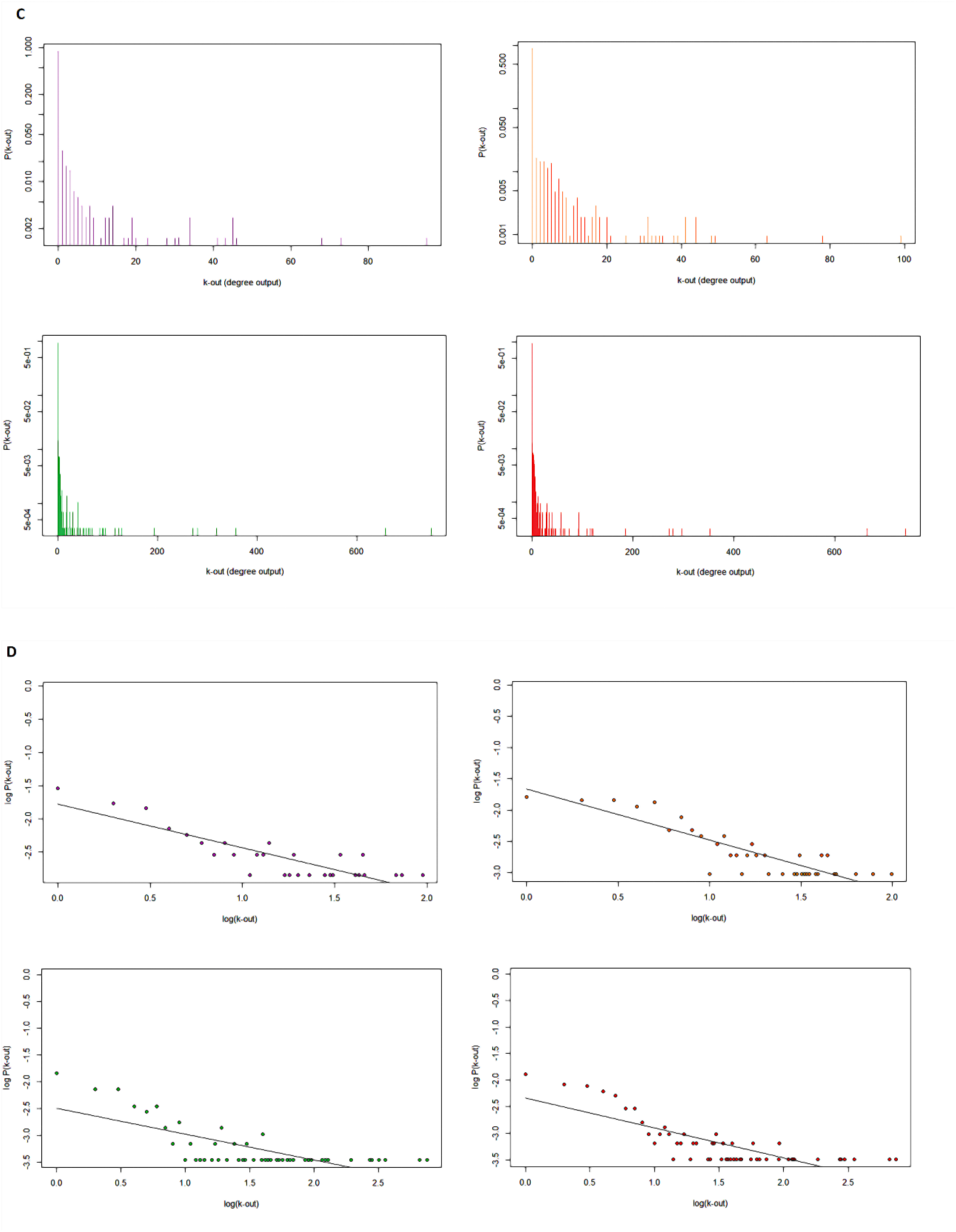
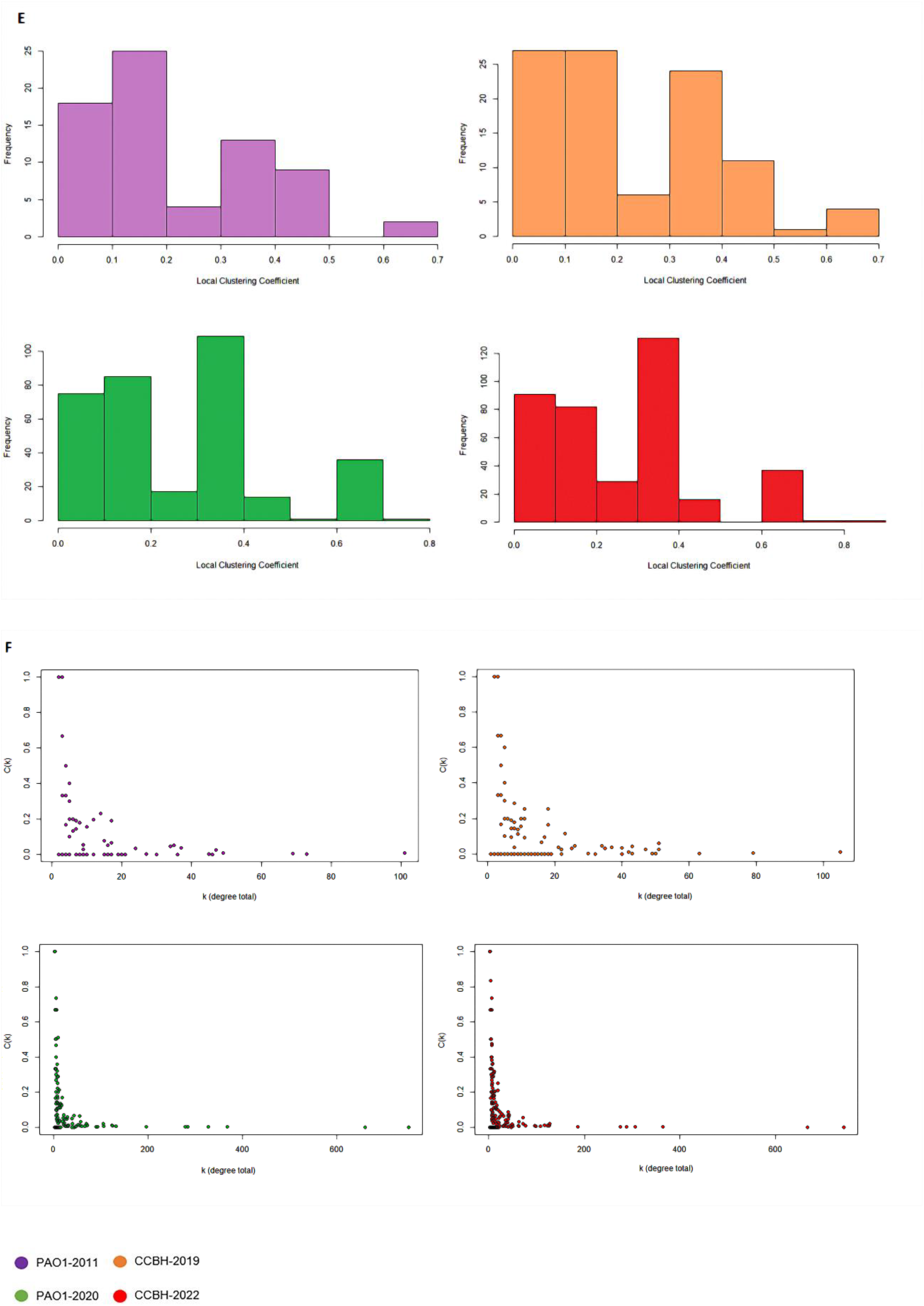
graphical representation of structural measurements of the CCBH-2022 (red) compared to the previously published networks: PAO1-2011 (purple), CCBH-2019 (orange), and PAO1-2020 (green). **A, B**: incoming degree distribution of the four GRNs; **C, D**: outgoing distribution of the four GRNs. The distributions are plotted on a linear (A, C) and on a logarithmic scale (B, D); **E**: local clustering coefficient distribution; **F**: clustering coefficient by degree.

The distribution of local clustering coefficients can be seen in Fig. 2E. The CCBH-2022 had a global clustering coefficient equal to 4.27e-03, higher than previous networks (PAO1-2020: 3.03e-03; CCBH-2019: 3.2e-02; PAO1-2011: 2.28e-02).

The scatter plot in Fig. 2F shows the correlation between the local clustering coefficient C(i) and the degree k(i).

Similar to the previous GRN, CCBH-2022 was disconnected, showing one large connected component (3034 genes) and 26 small connected components.

The most frequent mode of regulation in CCBH-2022 is activation, being 70,3% of the total interactions in the network, followed by 12% of repression mode and 17.6% of dual or unknown mode. Autoregulation occurs when a gene regulates its own expression, and the prevalence in the CCBH-2022 is of negative autoregulatory motifs.

The most abundant motif in all four networks was the coherent type I FFL, with 240 in the CCBH-2022 (PAO1-2011: 82; CCBH-2019: 79; PAO1-2020: 226). In addition, there were ten incoherent type II FFL motifs in the CCBH-2022 (PAO1-2011: 3; CCBH-2019: 4; PAO1-2020: 8).

Table II shows the 30 most influential hubs in the CCBH-2022.

**TABLE II.**
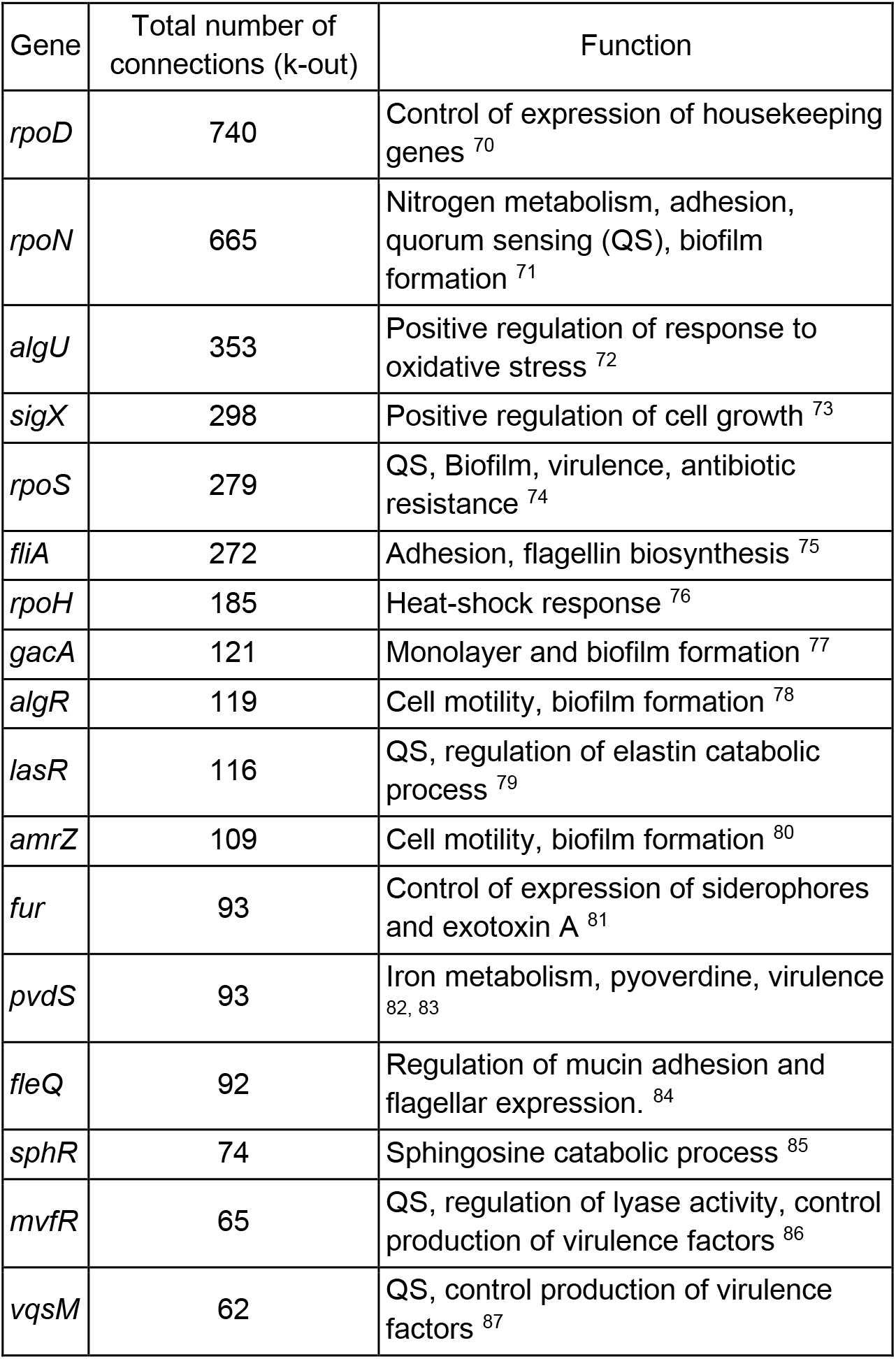
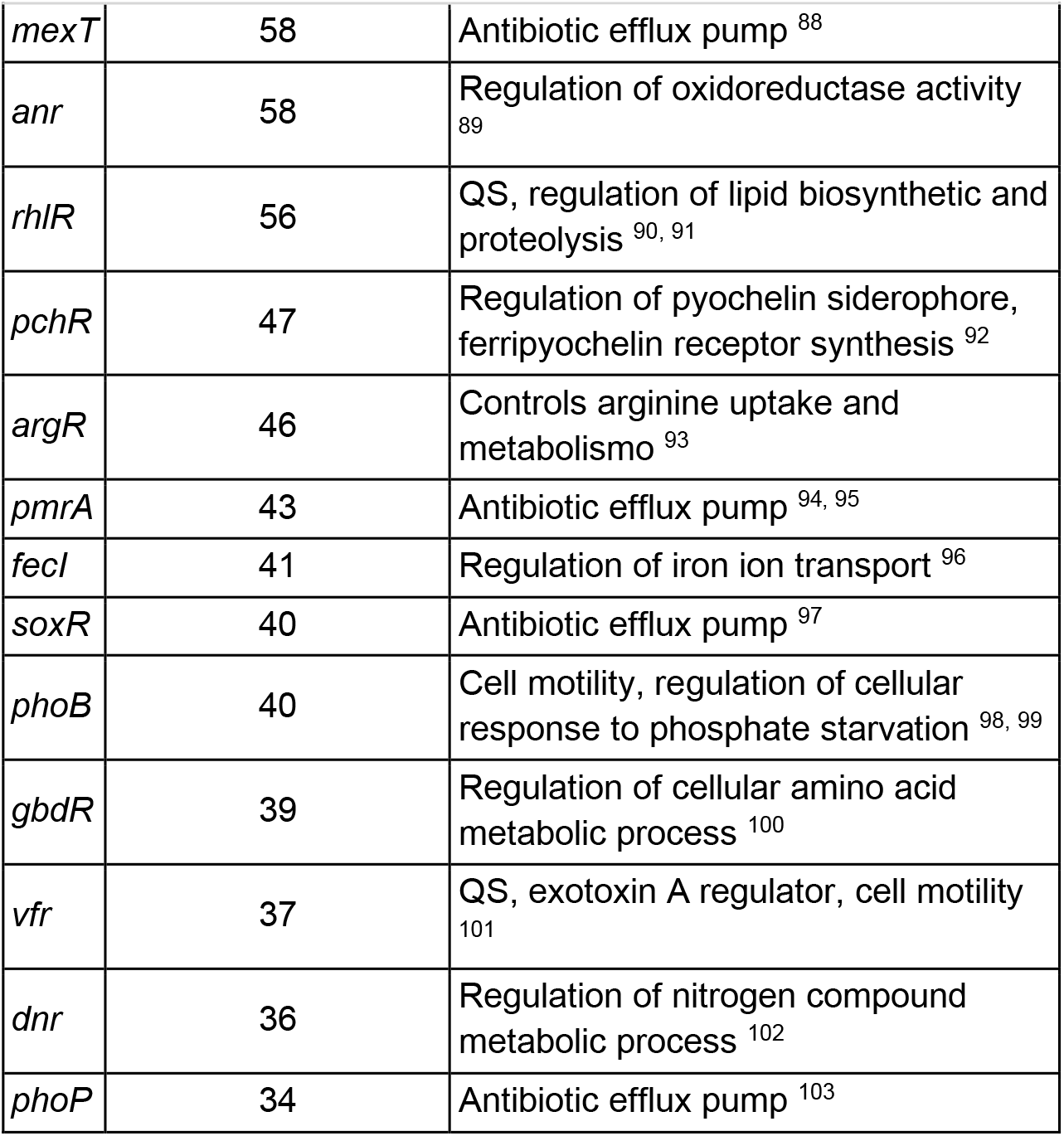
The 30 most influential hubs of the CCBH-2022

An analysis was performed to determine whether the hubs are interconnected through direct interactions (Fig.3).

## DISCUSSION

As one of the most important networks of bacteria described in the literature, the GRN of *P. aeruginosa* reconstruction and analysis contributes to giving a better understanding of its antibiotic resistance and for the knowledge of related cellular processes, such as adaptive and pathogenic, mainly when based on an MDR strain as CCBH4851.

In this work, we have good coverage of roughly 50% of the genome on this updated network. The genome of reference strain PAO1 has 6.2Mbp, and the PAO1-2020 has a coverage of 50% as well, with 5040 interactions and 3006 genes.^(20)^ However, considering that the CCBH4851 genome has 6.8Mbp and has 5507 edges, and 3291 nodes, we can affirm that, to the best of our knowledge, this study presents the largest GRN of *P. aeruginosa* that has been assembled to date.

On the structural aspects, the charts in Fig.2 and data in Table I make clear that CCBH-2022 represents a substantial improvement in terms of network completeness and complexity when compared with the previous *P. aeruginosa* GRNs, since it includes more nodes, edges, and network motifs, and when comparing clustering coefficients (Fig. 2E, F). For the *in silico* approach, the network structural analysis is essential to understand the network architecture and performance.

The structural measures of the CCBH-2022, such as node degree distribution and clustering coefficient, are consistent with a qualitative description of a scale-free network type. Indeed, the degree distribution followed the power-law distribution (Fig. 2B, D): a small number of nodes had many connections (the hubs) and many nodes had few connections.

The local clustering coefficient and node degree correlation (Fig. 2F) showed that nodes with lower degrees had greater local clustering coefficients than nodes with higher degrees. These characteristics are representative of several biological processes, e.g., RNA binding.^(104,105)^

The CCBH-2022 showed a lower density value than PAO1-2020. The density of both GRNs was low due to the dynamic and structural flexibility of the networks, a characteristic typical of natural phenomena-based networks (106), and because the nodes were not all interconnected.^(2)^ However, CCBH-2022 density was lower probably because it has 26 small connected components disconnected from the larger one (Fig.1), while the PAO1-2020 had 12 separated components. The variation in the number of connected components is plausible due to their size difference and the biological information about interactions available for the reconstruction.

All the previous *P. aeruginosa* GRNs are disconnected, showing one large connected component and a separated few small connected components, and there may be several reasons for this disconnection in specific points. According to Medeiros *et al*.^(2)^, interactions among all genes are not really expected since some genes in an organism are independent of each other, compartmentalized or global, constitutive or growth phase-dependent, and are triggered in different growth phases, thus resulting in a disconnected network, which corroborates with the observed low density. The reason can also be from loss of existing interactions or a gain of interactions still not fully described from additional strain-specific blocks of genes acquired by horizontal gene transfer.^(107)^

The large number of connected components found in the CCBH-2022 GRN results from connectivity parameters and the global clustering coefficient. Both structural measures are affected by the same biological behaviors.^(106)^

The most frequent regulatory activity in CCBH-2022 is activation, but ∼50% of the autoregulation was negative, which may be a consequence of the increase in negative regulation in the overall network interactions compared to the previous ones. Negative auto-regulation in biological systems is commonly observed.^(108)^ The *Escherichia coli* GRN exhibited the same pattern, with the negative autoregulation prevailing concurrently with the positive regulation in the overall network.^(109)^ The continuity of biological processes is ensured by positive autoregulation.^(110)^ For example, quorum sensing, biofilm formation, secretion of toxins, virulence and resistance factors production, once initiated, must reach a final stage in order to have the expected effect.^(2)^ In the CCBH-2022 GRN, genes involved in these processes, such as *lasR*,^(79)^ *rlhR*,^(91)^ *pvdS*,^(83)^ *algU*,^(72)^ *dnr*^(102)^ and *anr*,^(89)^ have positive autoregulation (and are hubs).

Negative cycles are also crucial for life-sustaining cyclic processes such as metabolic processes ^(111)^ and cellular homeostasis.^(112)^ In the CCBH-2022 GRN, genes involved in arginine metabolism (*iscR, desT, lexA, hutC*, and *mvat*)^(109)^ showed a predominance of negative mode of autoregulation. Negative autoregulation is associated with cellular stability.^(113)^ It provides a rapid response to variations in concentrations of proteins, toxins and (or) metabolites, to avoid undesired effects such as energy cost of unneeded synthesis.^(114)^ In the CCBH-2022 GRN, *algZ* (transcriptional activator of AlgD, involved in alginate production),^(115)^ *lexA* (involved in the SOS response),^(116)^ *metR* (involved in swarming motility and methionine synthesis),^(117,118)^ *ptxR* (affects exotoxin A production)^(119)^ and *rsaL* (quorum-sensing repressor)^(120)^ presented negative autoregulatory interactions. Autoregulation is common among genes positioned upstream in GRN with crucial developmental functions.^(121,122)^

The FFL motifs are essential for the modulation of cellular processes according to environmental conditions.^(123)^ CCBH-2022 has 951 FFL motifs, which are patterns of structural structures, while the PAO1-2020 has 702. There are 240 coherent type I FFL motifs in the CCBH-2022, an abundant presence. According to Mangan & Alon (2003)^(124)^ these motifs act as sign-sensitive delay elements i.e., a circuit that responds rapidly to steplike stimuli in one direction (ON to OFF), and as a delay to steps in the opposite direction (OFF to ON); the temporary removal of the stimulus ceases transcription, so the activation of expression requires a persistent signal to carry on.

The incoherent type II FFL motif was less represented but also found in all the GRN, with a total of 10 in the CCBH-2022 GRN. Contrastingly with the coherent FFL, the type II FFL acts as a sign-sensitive accelerator, i.e., a circuit that responds rapidly to step-like stimuli in one direction but not in the other direction.^(124)^

One last characteristic revealed by the structural analysis was the presence of hubs. The hub’s network (Fig. 3) shows the connection among their interactions; they are all interconnected, and belong to the largest connected component of the GRN (Fig.1A).

**Fig. 3:**
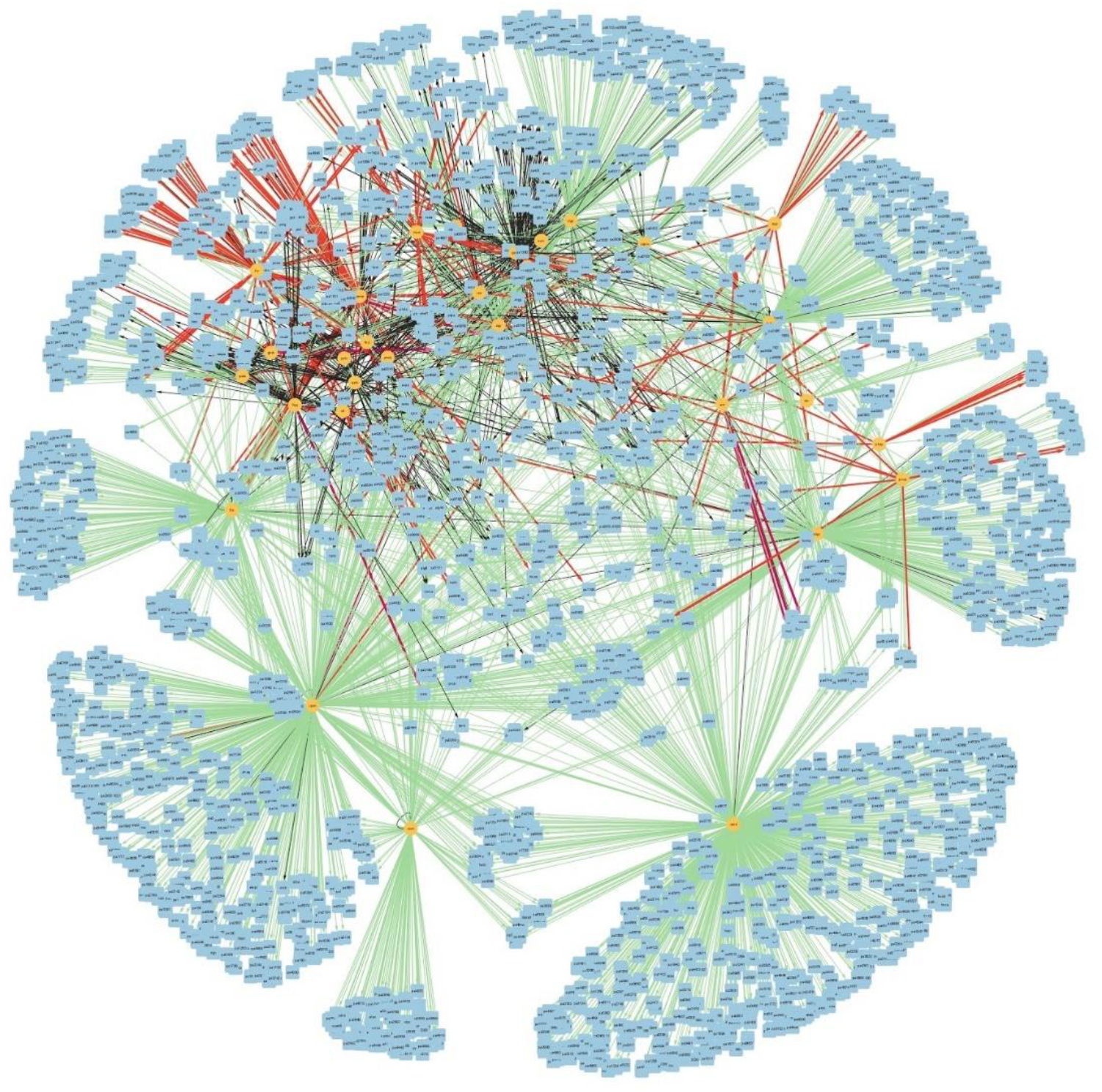
connectivity relationships among the 30 most influential hubs of the CCBH-2022. Yellow circles indicate regulatory genes considered hubs, light blue circles indicate target genes, black lines indicate an unknown mode of regulation, green lines indicate activation, and red lines indicate repression. Purple lines indicate a dual-mode of regulation.

This connectivity reflects the importance of the influential genes. The hubs can be considered the basis of the GRN. They are crucial in searching for potential drug targets for developing new drugs, as in a direct interaction with their specific targets or as for an indirect interaction with the subsequent process regulation triggered by them. The CCBH-2022 hubs are mainly associated with efflux pump mechanisms (*mexT, pmrA, soxR, phoP*),^(88,94,97,103)^ alginate biosynthesis (*algU, algR, rpoN*),^(125)^ and biofilm formation (*rpoN, rpoS, gacA, amrZ*).^(126)^ Table III shows the 30 hubs of PAO1-2020.

**TABLE III.** The 30 most influential hubs in the PAO1-2020

| Gene | Total number of connections (k-out) |
| --- | --- |
| *rpoD* | 749 |
| *rpoN* | 658 |
| *algU* | 357 |
| *sigX* | 319 |
| *fliA* | 281 |
| *rpoS* | 271 |
| *rpoH* | 194 |
| *gacA* | 128 |
| *algR* | 122 |
| *amrZ* | 115 |
| *lasR* | 95 |
| *pvdS* | 91 |
| *sphR* | 90 |
| *fleQ* | 85 |
| *fur* | 69 |
| *vqsM* | 65 |
| *mvfR* | 62 |
| *pchR* | 57 |
| *anr* | 53 |
| *mexT* | 51 |
| *argR* | 46 |
| *fecl* | 44 |
| *gbdR* | 42 |
| *rhlR* | 40 |
| *phoB* | 40 |
| *pmrA* | 40 |
| *soxR* | 39 |
| *dnr* | 34 |
| *himA* | 30 |
| *himD* | 30 |

They are overly similar to the CCBH-2022 hubs, with some changes in the k-out connections. However, two CCBH-2022 hubs were not present in PAO1-2020: *vrf*, a global virulence factor regulator ^(127)^ that directly regulates 37 genes, and *phoP*, associated with multidrug-resistance, including polymyxin,^(128,129)^ which directly regulates 34 genes. These facts strongly indicate that the operation of the main network hubs is not identical. The functioning of CCBH4851 is different given the presence of these 2 critical genes associated with multi-drug resistance mechanisms.

The *Vfr* gene’s role in regulation of virulence factor production is related to the production of exotoxin A, a toxin that modifies specific target proteins within mammalian cells and induces necrosis in different tissues and organs in MDR *P. aeruginosa* infections.^(130,131)^ The *Vfr* gene also regulates the *las* and consequently the *rhl* quorum-sensing system, two systems that together control expression of several genes associated with virulence factor production,^(132)^ including alkaline protease, exotoxin A, pyocyanin, and rhamnolipid, as well as critical genes such as *rpoS* (the 5th most influential gene in CCBH-2022).^(133)^ The signal receptor (*R* gene) is one of the essential components of the *las* and *rhl* QS systems. It is necessary for coding the transcriptional activator protein (R protein).^(134)^ The *lasR* and *rhlR* genes are among the 20 principal hubs. The *phoP* gene has an essential role in MDR *P. aeruginosa*, being involved in more than one drug-resistant mechanism. Apart from their role in two-component regulatory systems with *phoQ* gene, it also plays significant roles in multidrug efflux pumps and alteration in drug targets.^(135)^ Miller *et al*. (2011)^(128)^ showed that deletion of the phoPQ two-component regulatory system locus in a laboratory-adapted *P. aeruginosa* PAK, which is considered an MDR strain, resulted in loss of polymyxin resistance.^(136)^

*P. aeruginosa* evades the antimicrobial activity during treatment and exerts antimicrobial resistance by mainly intrinsic resistance mechanisms. Examples of resistance mechanisms are multi-drug efflux pumps, biofilm synthesis, enzymatic inactivation/degradation, drug permeability restriction, production of beta-lactamases, acquired resistance by a mutation in drug targets, and acquisition of resistance genes via horizontal gene transfer.^(135)^

There is a directed regulatory connection from alginate biosynthesis to iron metabolism and some antibiotic resistance mechanisms.^(137)^ The *algU, algR, rpoN, pvdS* and *fecI* genes are related to these processes^(138,139)^ and are among the most influential hubs.

*P. aeruginosa* has multiple efflux pump systems that prevent the antimicrobial agents from accumulating in adequate concentration to cause an effect in the cell, extruding the drug out.^(135)^ Efflux pump systems are associated with resistance to beta-lactams, fluoroquinolones, tetracycline, chloramphenicol, macrolides, and aminoglycosides.^(140)^ Differential expression or mutations of efflux system genes are also contributing factors for both carbapenem and aminoglycoside resistance.^(141)^ The *mexT, pmrA, soxR* genes, related to multidrug antibiotic efflux pumps, are also amongst the most influential hubs. The *fleQ* gene is also among the hubs and affects *psl* (polysaccharide synthesis locus) genes and the regulation of the efflux pump genes, *mexA, mexE*, and *oprH*, by *brlR*.^(2,142)^ The *psl* cluster comprises 15 exopolysaccharide biosynthesis-related genes organized in tandem that are important for biofilm formation.^(143)^ The *mexT* and *soxR* genes positively regulate an efflux pump system and several virulence factors,^(144,145)^ and *pmrA* regulates efflux pumps and the polymyxin B and colistin resistance.^(95,146,147)^

Efflux pumps also help biofilm formation.^(148)^ Biofilms are also related to protection from the host immune system and antibiotic penetration and tolerance, preventing them from entering the microbial population, inhibiting its action as a first-line defense mechanism.^(123,149,150)^

The *rpoN, rpoS, gacA, algR* and *amrZ* hubs participate in the regulation of *P. aeruginosa* biofilm.

This system biology approach to characterize the MDR *P. aeruginosa* CCBH4851 regulatory network may lead to the development of strategies to disrupt the connectivity of these essential processes, thus, possibly decreasing the pathogenicity and suppressing the resistance of this bacterium.

## CONCLUSIONS

This manuscript reports the reconstruction and structural analysis of the largest *P. aeruginosa* regulatory network to date.

This work can give new insights into identifying novel candidate antibiotic targets and contributes to an increase in our understanding of the behavior of this bacterium.

This network’s dynamic model construction is one of our future studies, intending to help researchers working on experimental drug design and screening. The goal is to predict the dynamic behavior better and improve the understanding of *P. aeruginosa*, allowing the simulation of normal and stress conditions to discover potential therapeutic targets and help develop new drugs against *P. aeruginosa’s* bacterial infection.

## ACKNOWLEDGEMENTS

To Inova-Fiocruz (grant #VPPCB-007-FIO-18-2-117), Faperj and Capes, for financial support.

## AUTHOR’S CONTRIBUTIONS

MSC performed the GRN reconstruction, its visualization and drafted the manuscript. FMF performed the structural analysis. FABS supervised this study. FMF, MTS, MAM, APDCA and FABS provided scientific advice and contributed to revision of the text. All authors read and approved the final manuscript.

